# Dynamics of MicroRNA Secreted via Extracellular Vesicles During the Maturation of Embryonic Stem Cell-Derived Retinal Pigment Epithelium

**DOI:** 10.1101/2024.03.31.587478

**Authors:** Dimitrios Pollalis, Gopa Kumar Gopinadhan Nair, Justin Leung, Clarisa Marie Bloemhof, Jeffrey K. Bailey, Britney O. Pennington, Kaitlin R. Kelly, Amir I. Khan, Ashley K. Yeh, Kartik S. Sundaram, Dennis O. Clegg, Chen-Ching Peng, Liya Xu, Constantin Georgescu, Jonathan Wren, Sun Young Lee

## Abstract

Retinal pigment epithelium (RPE) cells are exclusive to the retina, critically multifunctional in maintaining the visual functions and health of photoreceptors and the retina. Despite their vital functions throughout lifetime, RPE cells lack regenerative capacity, rendering them vulnerable and central to degenerative retinal diseases. With advancements in stem cell technology enabling the differentiation of functional cells from pluripotent stem cells and leveraging the robust autocrine and paracrine functions of RPE cells, extracellular vesicles (EVs) secreted by RPE cells hold significant therapeutic potential in supplementing RPE cell activity. While previous research has primarily focused on the trophic factors secreted by RPE cells, there is a lack of studies investigating miRNA, which serves as a master regulator of gene expression. Profiling and defining the functional role of miRNA contained within RPE-secreted EVs is critical as it constitutes a necessary step in identifying the optimal phenotype of the EV secreting cell and understanding biological cargo of EVs to develop EV-based therapeutics. In this study, we present a comprehensive profile of miRNA in small extracellular vesicles (sEV) secreted during RPE maturation following differentiation from human embryonic stem cells (hESCs). This exploration is essential for ongoing efforts to develop and optimize EV-based intraocular therapeutics utilizing RPE-secreted EVs, which may significantly impact the function of dysfunctional RPE cells.

## Introduction

Retina pigment epithelium (RPE) is a monolayer of polarized cells uniquely located between the photoreceptors and the choroid vasculature within the chorioretina. These cells serve highly specialized functions in maintaining the health and visual functions of photoreceptors and the retina, including conservation of visual cycle, protection against oxidative stresses, secretion of cytokines and growth factors, maintain blood retinal barrier (BRB) and transportation of nutrients (Strauss, 2005). Given their postmitotic status and the demanding roles they play throughout the lifetime, the RPE is particularly susceptible to pathology in major blinding retinal degenerative conditions such as age-related macular degeneration (AMD) (Handa et al., 2019; Campello et al., 2021).

Benefiting from the advancements in stem cell technology enabling the differentiation of functional cells from pluripotent stem cells (PSC) or induced PSC (iPSC) and leveraging the robust autocrine and paracrine functions of RPE cells, extracellular vesicles (EVs) secreted from RPE cells hold significant therapeutic potential in supplementing RPE cell activity. Studies have shown that stem cell-derived polarized RPE cells exhibit both phenotypic and functional similarities to RPE cells obtained from human retina tissues (Croze et al., 2014; Leach et al., 2016). These fully differentiated and polarized RPE cells have demonstrated proven intraocular safety profiles and functional integration in ongoing human clinical trials involving cell transplantation (Schwartz et al., 2015; Mehat et al., 2018; Mandai et al., 2017; Kashani et al., 2018; Kashani et al., 2021). Our group recently reported that concentrated cell culture conditioned media (CCM) of hESC-derived fully differentiated and polarized RPE promoted retinal progenitor cell survival, reduced oxidative stress in APRE-19 cells, and preserved PR and their function in the Royal College of Surgeons (RCS) rat model (Ahluwalia et al., 2023). Accumulating evidence suggests that cell secreted small extracellular vesicles (sEV) play a crucial role in facilitating cell-to-cell communication. sEV carries essential biological cargos such as proteins, lipids, and microRNAs (miRNA), which are encapsulated and protected by a lipid bilayer membrane of sEV from nucleases, proteases, changes in pH or osmolarity (Ratajczak et al., 2006; van der Pol et al., 2012; Ludwig and Giebel, 2012). Taking advantage of advancements in high-throughput technologies, such as proteomics, microarray, RNA-Seq, and ChIP-Seq, systematic analysis of EV cargo has been enabled (Kim et al., 2017; Vaka et al., 2023). However, studies on miRNA secreted from RPE are lacking, especially during their maturation followed by differentiation from PSC while RPE-secreted proteomics have been relatively well defined (Meyer et al., 2019; Flores-Bellver et al., 2021; Yuan et al., 2015; La Torre et al., 2013; Georgi and Reh, 2010; Wright et al., 2020).

miRNAs are small (∼22 nucleotides) noncoding, highly conserved, single-stranded RNAs known to play an important role in various biological processes by directly or indirectly regulating the expression of target mRNAs (Carthew and Sontheimer, 2009; Kim, 2005). Approximately 2200 miRNA genes have been reported to exist in the mammalian genome and the expression of miRNAs is tightly regulated, showing dependence on developmental stage and tissue specificity (Ardekani and Naeini, 2010; Lagos-Quintana et al., 2002). Furthermore, their associated with retinal health and diseases have been demonstrated (Yuan et al., 2015; La Torre et al., 2013; Georgi and Reh, 2010; Wright et al., 2020). Thus, the miRNA cargo of EVs represents a potentially critical component of the functional role of RPE-secreted EVs in therapeutics. Profiling and understanding the functional role of miRNA contained within RPE-secreted EVs are critical steps in identifying the optimal phenotype of the EV secreting cell and understanding biological cargo of EVs to develop EV-based therapeutics.

RPE maturity has been reported to affect cellular morphology [27,28], pigmentation [,32] and the production of several functional proteins such as pigment epithelium-derived factor (PEDF), bestrophin-1 (BEST1) and cellular retinaldehyde-binding protein (CRALBP) [20] (Dunn et al., 1996; German et al., 2008; Bennis et al., 2007). Previous studies have demonstrated that in vitro maturation of hESC-derived RPE cells continues for 8 weeks with increasing expression of key functional genes. Interestingly, transplantation of hESC-derived RPE obtained after 4 weeks of culture showed a better vision-rescuing effect in animal models compared to those cultured for 2 or 8 weeks (Al-Ani et al.,2020; Davis et al., 2017). This suggests that the optimal phenotype of the cells needs to be assessed based on the therapeutic strategy employed, such as cell implantation or cell secreted EVs.

In this study, we present a comprehensive miRNA profiling in sEV secreted during the maturation of RPE after differentiation from hESC; *early-stage* hESC-RPE (11-22 days in culture), *mid-stage* hESC-RPE (28-39 days in culture), and *late stage* hESC-RPE (59-70 days in culture). Additionally, we conducted associated pathway analyses to gain insights into the functional roles of these profiled miRNAs. Our findings contribute to bridging the existing knowledge gap and lay the foundation for further exploration of RPE-secreted EV therapeutics.

## Result

### hESC-RPE Characterization

The fully differentiated hESC-RPE cells, regardless of their maturation status, exhibited RPE cell characteristics, including distinctive ZO-1-positive cobblestone, hexagonal morphology. Meanwhile, the pigmentation grading increased with maturation status across three groups, *early-stage* (11-22 days in culture)*, mid-stage* (28-39 days in culture), and *late-stage* (59-70 days in culture). (**Figure 1A**). Additionally, immunofluorescence analysis of hESC-RPE cells, stained with anti-BEST1 and anti-RPE65 antibodies, revealed maturation-dependent variations. These differences were further confirmed through qPCR-based evaluation of gene expression, with BEST1 expression levels being 31.8-fold, 46.2-fold, and 28.9-fold increased for *early-stage*, *mid-stage*, and *late-stage* hESC-RPE, respectively, and RPE65 expression levels being 0.12-fold, 0.3-fold, and 0.28-fold increased for *early-stage*, *mid-stage*, and *late-stage* hESC-RPE, respectively (**Figure 1B and 1C**).

**Figure 1.**
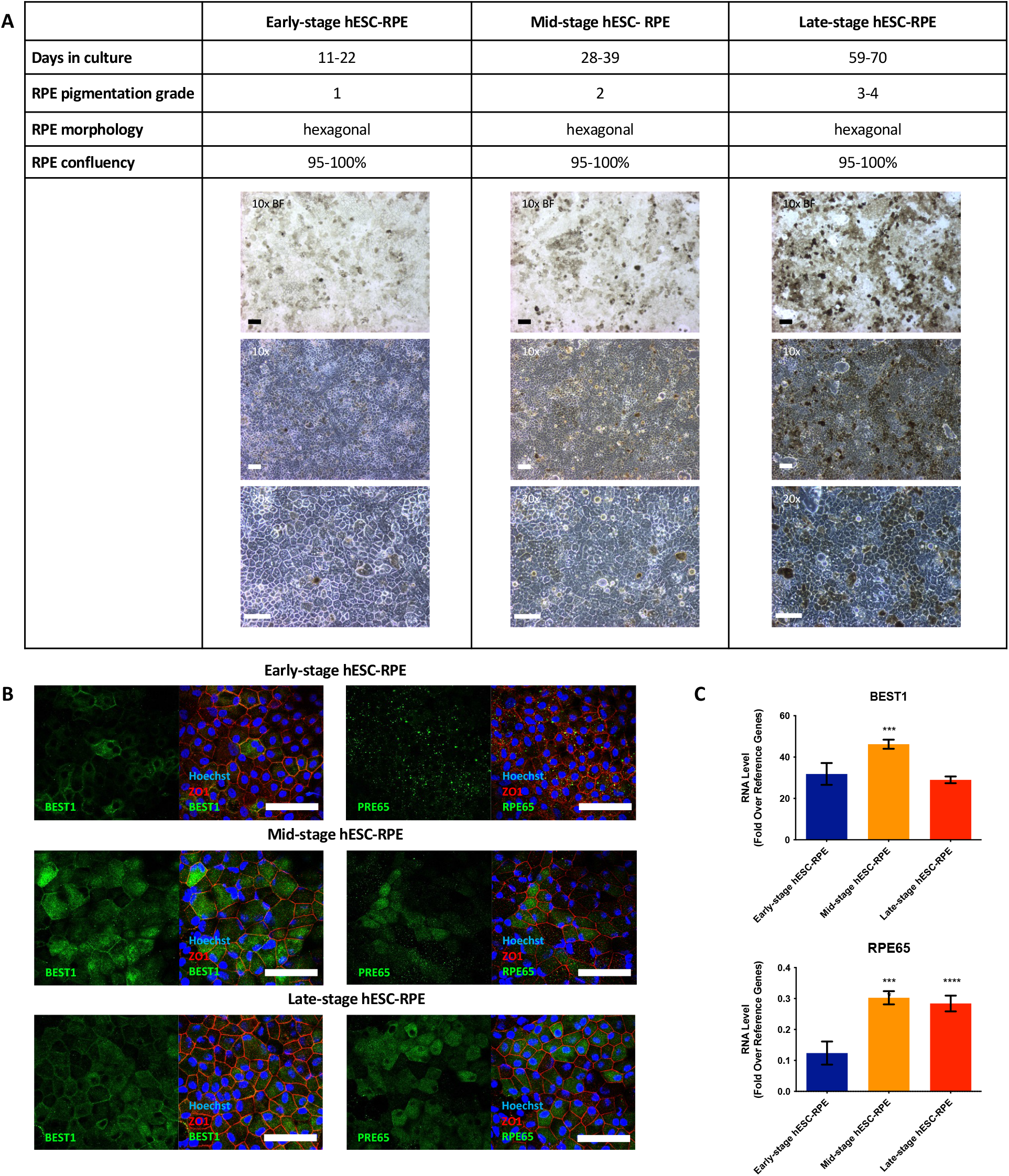
**A.** Summary table of the phenotypical characteristics of *early-stage* (11-22 days in culture), *mid-stage* (28-39 days in culture), and *late-stage* (59-70 days in culture) of maturation in hESC-RPE cells cultured, along with representative brightfield and phase-contrast images (scale bar, 100 μm). **B.** Representative images of hESC-RPE cells stained with anti-BEST1, RPE65, and ZO1 antibodies (scale bar, 50 μm). **C.** BEST1 expression was increased in *mid-stage* hESC-RPE compared to the other two groups, while RPE65 expression was found to be increased during the maturation of hESC-RPE cells. N=4, *** *p* < 0.001, **** *p* < 0.0001.

hESC-RPE cells presented fully differentiated distinctive RPE characteristics, with increased pigmentation observed over RPE cell maturation, along with a decrease in their proliferation in cell culture (data not shown), with days of gap between the groups to minimize potential molecular overlapping.

### hESC-RPE-sEV Biophysical Characteristics

The quantification and characterization of the hESC-RPE-sEV were performed using NTA with ZetaView. Notably, the particle numbers of sEV released by *mid-stage* hESC-RPE exceeded those of the other groups, with counts for *early-*, *mid-*, and *late-stage* hESC-RPE-sEV measured at 6.84 x 10^10^ (± 6.61 x 10^9^), 1.9 x 10^11^ (± 9.98 x 10^9^), and 7.42 x 10^10^ (± 8.04 x 10^9^) particles/mL, respectively. The average size of sEV was comparable with 113.9 nm, 111 nm, and 110.4 nm in *early-*, *mid-*, and *late-stage* hESC-RPE-sEV, respectively, and their morphology observed in TEM images from the three age groups was comparable (**Figure 2A**).

**Figure 2.**
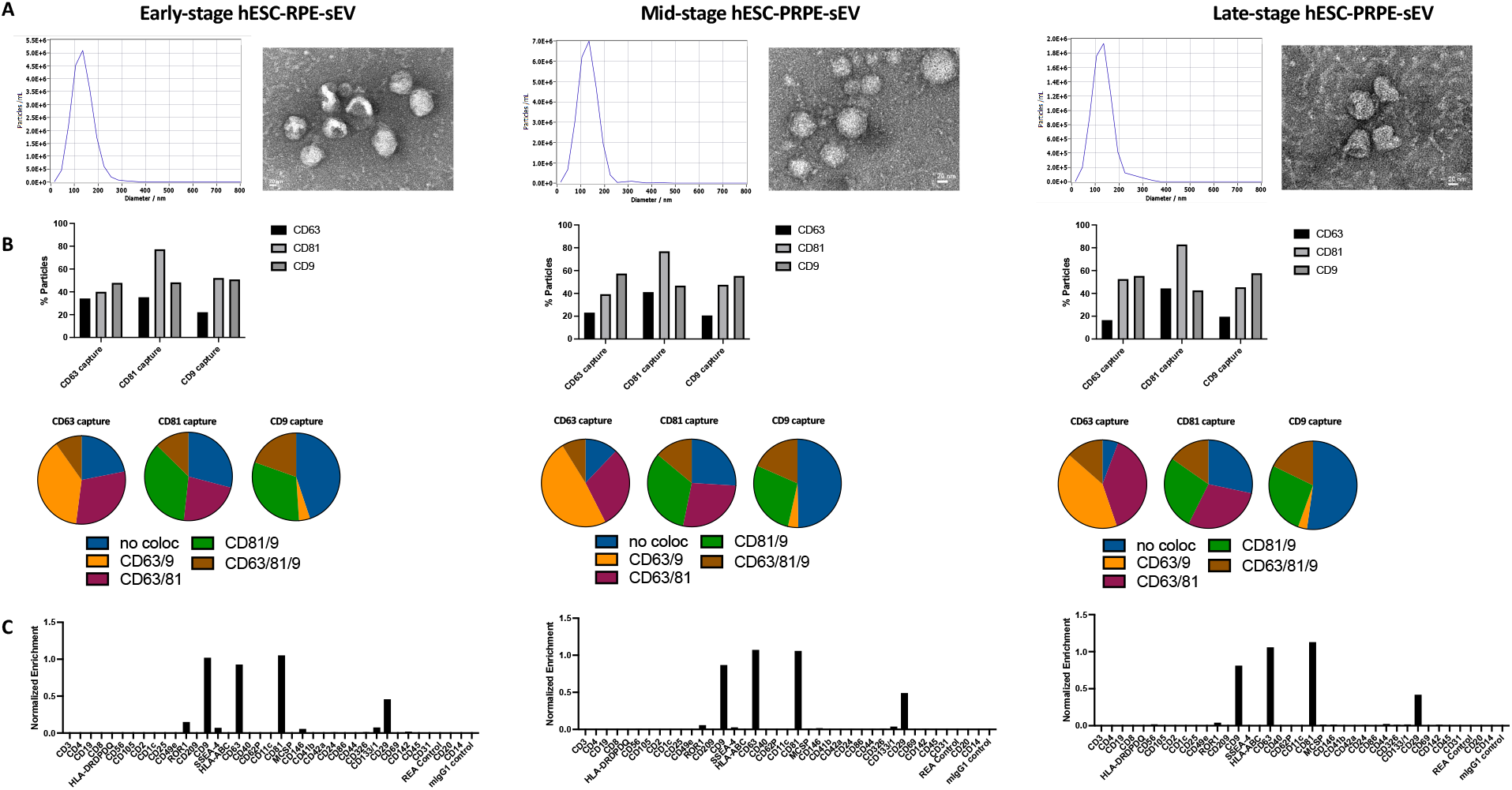
**A.** Representative size versus concentration distribution graphs of sEV derived from the three age groups of hESC-RPE cells as analyzed by ZetaView and representative transmission electron microscopy (TEM) images of sEV providing a visual comparison. **B.** Distribution and colocalization analysis of tetraspanins (CD81, CD9, and CD63) within sEV derived from the three age groups of hESC-RPE cells. **C.** Results from the MACSPlex assay conducted on *early-stage* (11-22 days in culture)*, mid-stage* (28-39 days in culture), and *late-stage* (59-70 days in culture) hESC-RPE-sEV.

### Exosomal Tetraspanins and Surface Epitopes of hESC-RPE-sEV

The tetraspanin distribution in sEV, as analyzed by ExoView, remained consistently similar across different maturation stages of the hESC-RPE cells (**Figure 2B**). In-depth surface protein epitopes capturing 37 distinct surface markers identified predominant surface proteins across different age groups, including ROR1, CD9, CD63, CD81, CD133/1, and CD29 (**Figure 2C**). Notably, the uniform tetraspanin distribution in hESC-RPE-sEV, combined with the extended surface epitope profile, differed from that of sEV isolated from HEK293T cells, a non-ocular reference cell line (**Supplementary Figure 1**). This suggests that our sEV characterization is sensitive enough to differentiate sEV derived from different cell sources.

### Purity, Protein, and RNA Concentration in hESC-RPE-sEV

The protein concentration was highest in the mid-stage hESC-RPE-sEV compared to *the early-* and *late-stage* groups. Further analysis of the sEV purity index (particle-to-protein ratio) indicated an increase in protein along with the elevated sEV counts, registering as 0.26, 0.84, and 0.26 mg/mL *in early-*, *mid-*, and *late-stage* hESC-RPE-sEV, respectively (**Figure 3A and 3B**). Therefore, the purity index (particle-to-protein ratio) of recovered sEV in all groups was found to be high and comparable among the groups (**Figure 3C**). The total RNA concentration also followed a similar pathway with comparable concentrations among the groups (8.488, 9.496, and 5.941 ng/μL in *early-*, *mid-*, and *late-stage* hESC-RPE-sEV, respectively) (**Figure 3D**).

**Figure 3.**
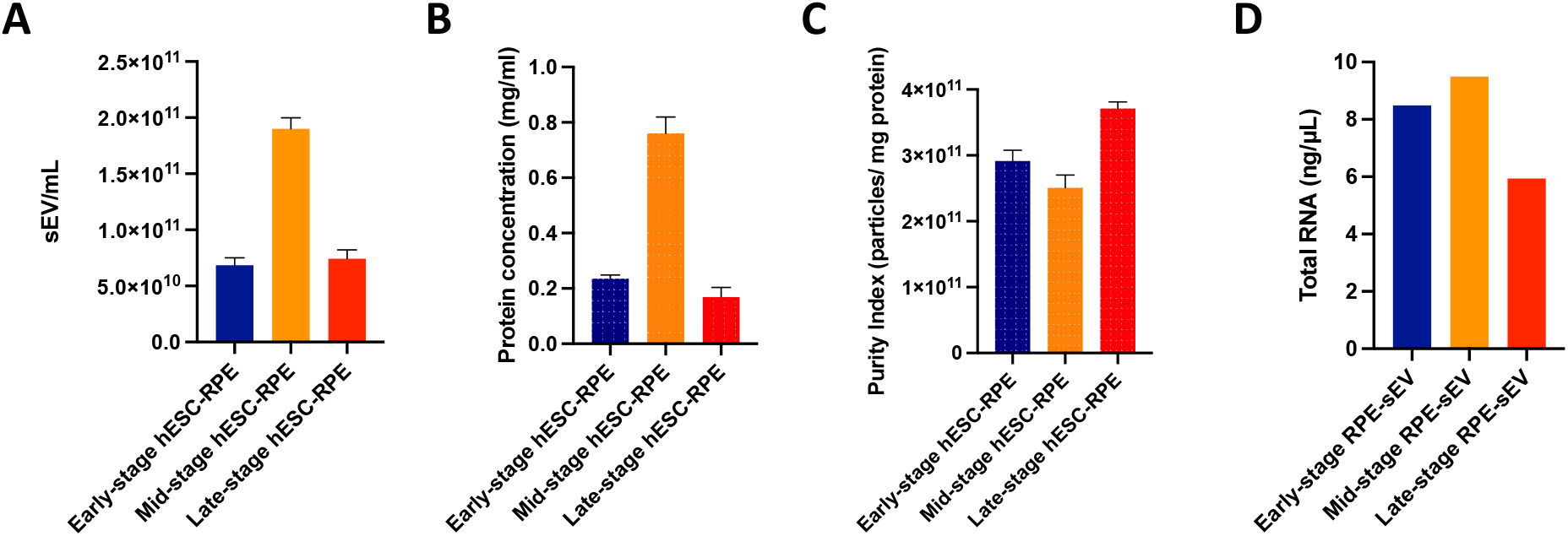
Particle numbers (**A**), protein concentration (**B**), purity index (particle-to-protein ratio) (**C**), and total RNA amount (**D**) from the recovered sEVs obtained from the same volume input of CCM, with similar cell culture numbers, were compared across the *early-stage* (11-22 days in culture)*, mid-stage* (28-39 days in culture), and *late-stage* (59-70 days in culture) hESC-RPE maturation.

### microRNA Profiles in sEV During hESC-RPE Maturation

The miRNA profiles of hESC-RPE-sEV exhibited notable distinctions across the three groups of hESC-RPE cells (**Figure 4A, Supplementary Tables 1, 2, and 3**). In the overall miRNA expression pool, comprising 94 miRNAs, varying counts were observed in sEV derived from *early-*, *mid-*, and *late-stage* hESC-RPE cells, with totals of 27, 85, and 34 miRNAs, respectively. Eleven miRNAs were shared only by early-stage and *mid-stage* hESC-RPE-sEV, while 15 miRNAs were shared exclusively between mid-stage and late-stage hESC-RPE-sEV. No miRNA was expressed solely in early- and *late-stage* hESC-RPE-sEV. *early-*, *mid-*, and *late-stage* hESC-RPE-sEV exhibited unique expression patterns for 3, 46, and 6 miRNAs, respectively. Thirteen miRNAs were identified as shared among all the hESC-RPE-sEV groups, with 9 of them (hsa-miR-720, hsa-miR-648, hsa-miR-627, hsa-miR-30c, hsa-miR-30b, hsa-miR-17, hsa-miR-1298, hsa-miR-1274B, hsa-miR-1274A) also observed in HEK293T-sEV (**Supplementary Figure 2** **Supplementary Tables 4**). A non-supervised hierarchical clustering of the 94 expressed miRNAs revealed a distinct separation of the mid-stage hESC-RPE-sEV group from the other two groups. (**Figure 4B**).

**Figure 4.**
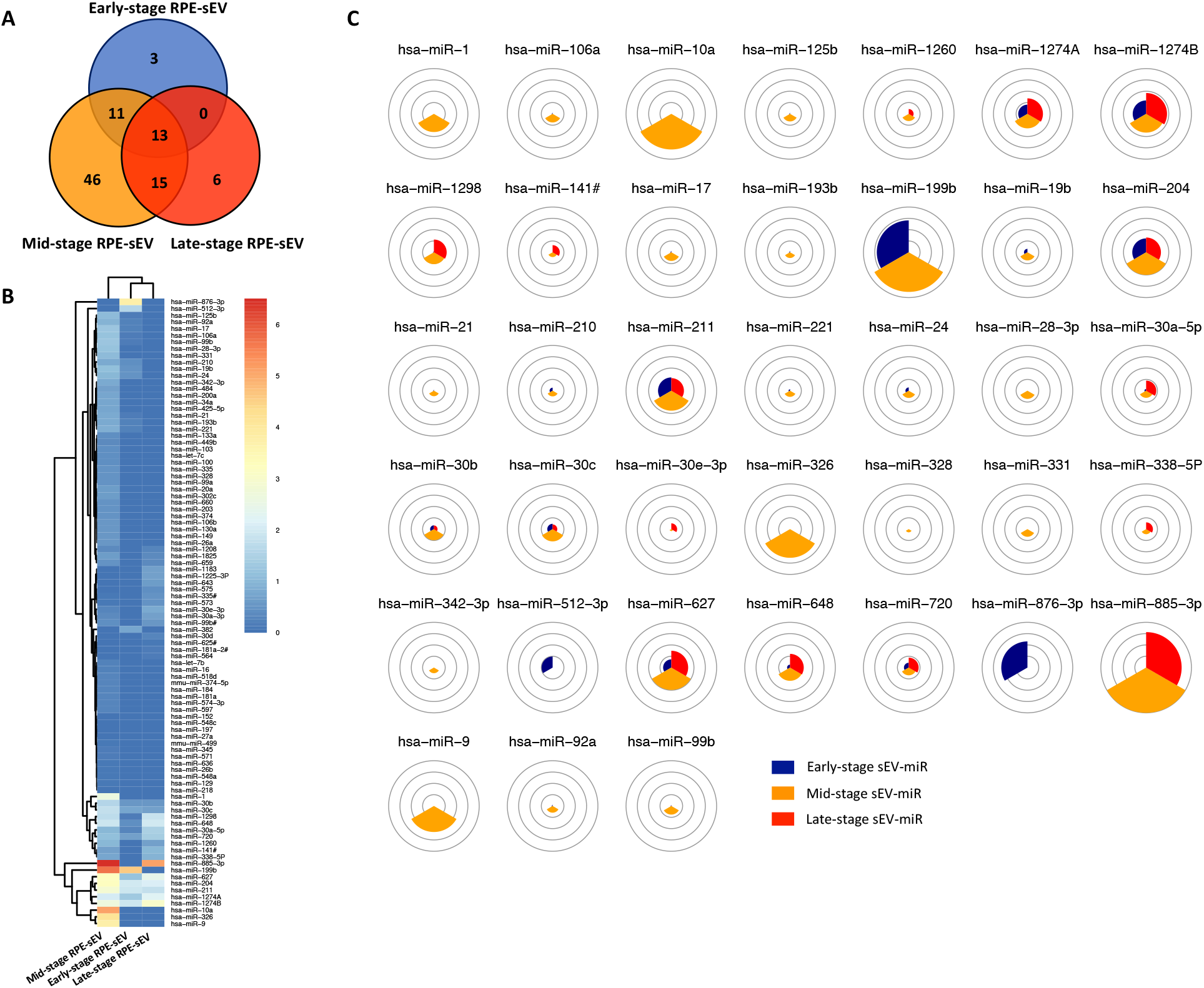
**A.** Venn diagram of the total number of 94 miRNAs identified (*Ct* values ≤ 30) contained in hESC-RPE-sEV across the *early-stage* (11-22 days in culture)*, mid-stage* (28-39 days in culture), and *late-stage* (59-70 days in culture) of hESC-RPE maturation, totaling 27, 85, and 34 miRNAs, respectively. **B.** Heatmap of the 94 expressed miRNAs in hESC-RPE-sEV throughout RPE maturation. **C.** Of 94 expressed miRNAs, top 5-fold differentially expressed miRNAs hESC-RPE-sEV during RPE maturation. miRNA expressions are presented as log2 transformed values.

### Pathway Analysis

To comprehensively investigate the functional implications of differentially expressed miRNAs across three distinct age groups, we conducted a gene set and pathway enrichment analysis focused on the top 5-fold expressed miRNA (**Figure 4C**). Using Ingenuity Pathway Analysis, we predicted target pathways for the top 5-fold miRNAs, identifying 511 statistically significant distinct pathways (**Supplementary Figure 3**). Our detailed examination, with a focus on extracellular vesicle- and retina-related pathways, revealed major associations with regulatory networks governing extracellular vesicle trafficking, tissue and cellular polarity, growth factor regulation, inflammation, and oxidative stress & senescence (**Figure 5**). Notably, pathways associated with major extracellular vesicle trafficking, such as clathrin-mediated endocytosis signaling and clathrin-associated endocytosis, were evident across all hESC-RPE-sEV groups. Among the pathways associated with tissue and cell architecture, the planar cell polarity (PCP) pathway emerged as a consistently regulated pathway by miRNAs secreted by hESC-RPE-sEV, irrespective of RPE age. Both *mid-* and *late-stage* hESC-RPE-sEV secreted miRNAs associated with growth factors responsible for tissue proliferation and repair, including BMP, HGF, and TGF-β (Kramer et al., 2021; Simó et al., 2006). Furthermore, miRNAs from mid-stage hESC-RPE-sEV influenced cell proliferation and tissue development-related pathways like EGF and FGF (Simó et al., 2006; Baker, 2001). Our analyses also indicated potential regulation of neurotrophic signaling pathways such as NGF and CNTF (Fischer et al., 2004). Additionally, VEGF and PDGF pathways crucial for vascular integrity in wound healing and tissue repair were predominantly associated with miRNAs derived from *mid-stage* hESC-RPE-sEV (Simó et al., 2006). Interestingly, the well-established PEDF regulation pathway, a major regulator for anti-angiogenesis and neuroprotection, exhibited regulatory associations with both *mid-* and *late-stage* hESC-RPE-sEV (Simó et al., 2006). Furthermore, miRNAs secreted via both *mid-* and *late-stage* hESC-RPE-sEV were implicated in the regulation of inflammation through various interleukin pathways. Finally, our study showed a significant association between predicted target genes and pathways related to oxidative stress and senescence.

**Figure 5.**
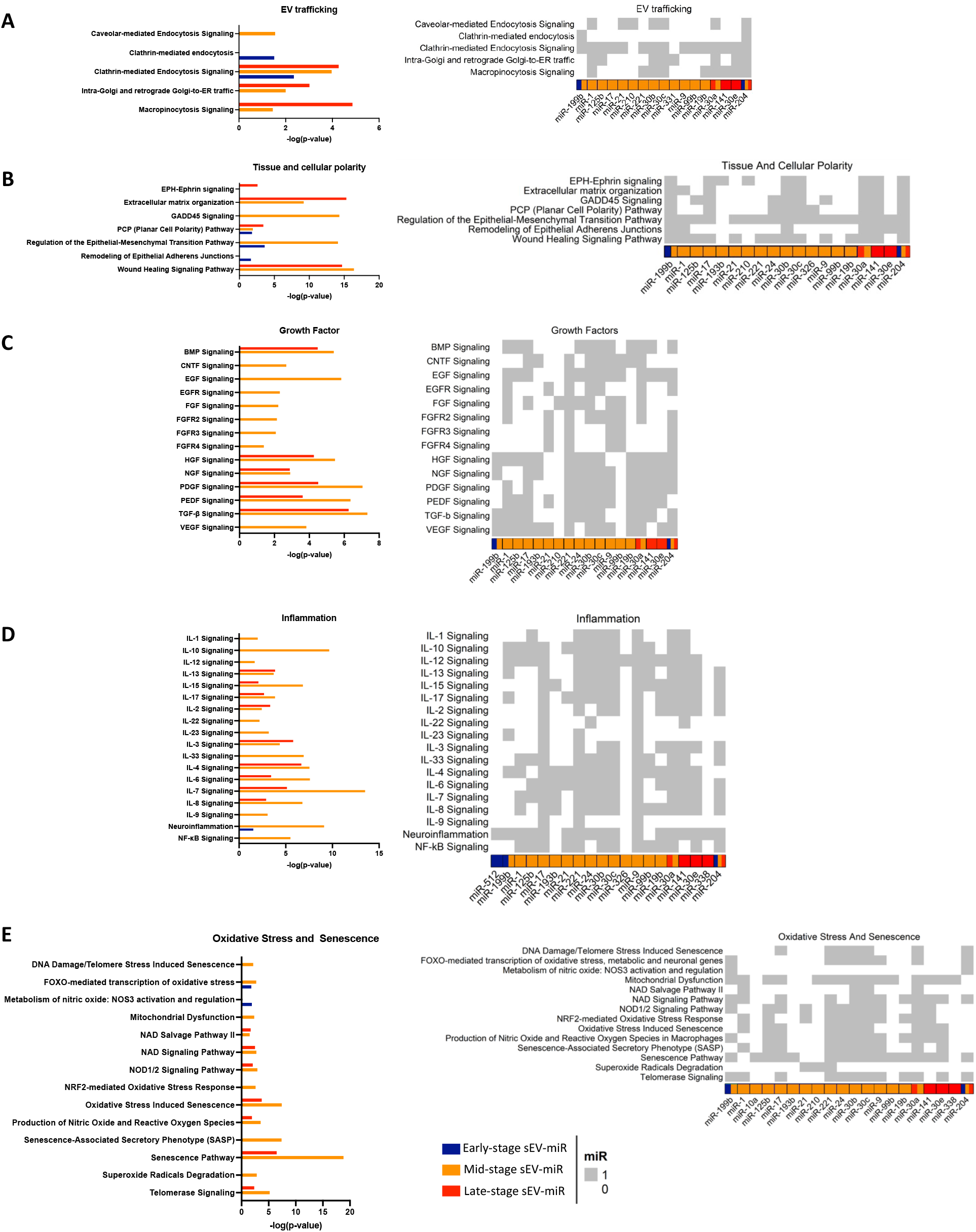
Of the total number of 94 miRNAs identified (with Ct values ≤ 30) contained in hESC-RPE-sEVs across the *early-stage* (11-22 days in culture)*, mid-stage* (28-39 days in culture), and *late-stage* (59-70 days in culture) of hESC-RPE maturation, totaling 27, 85, and 34 miRNAs, respectively, the top 5-fold expressed miRNAs were analyzed via Ingenuity Pathway Analysis (IPA) and IREDESCENT, focusing on pathways related to “extracellular vesicle” and “retina”. These pathways include (**A**) EV trafficking, (**B**) tissue/cellular polarity, (**C**) growth factors, (**D**) inflammation, and (**E**) oxidative stress & senescence. -Log (*p* value) > 1.3 is considered statistically significant.

### Pigment Epithelium-Derived Factor

A gradual increase in PEDF expression was observed with days in culture, with fold changes of 195.4 in early-stage, 277.4 in mid-stage, and 298.3 in *late-stage* hESC-RPE cells (**Figure 6A**). To further evaluate the secretory function of PEDF from hESC-RPE cells across different groups, we performed enzyme-linked immunosorbent assay (ELISA) on their conditioned CCM (**Figure 6B**). Additionally, we quantified the PEDF concentration in two distinct fractions of the hESC-RPE secretome: sEV-fraction and sEV-depleted (CCM minus sEV). Our findings revealed a 60% increase in PEDF secretion by both *mid-*and *late-stage* hESC-RPE cells compared to the *ealy-*stage group. In the sEV-depleted fraction (CCM minus sEV), PEDF concentration remained comparable across all age groups. In contrast, the sEV-enriched fraction of the conditioned CCM exhibited the highest concentration of PEDF in the mid-stage hESC-RPE group, supporting the notion that a portion of PEDF secreted by RPE is bound to EVs. Focusing on PEDF, we constructed and visualized networks that explore age-dependent miRNA-gene target interactions using Cytoscape. Pathway analysis of miR-gene target interactions revealed numerous miRNA associations (**Figure 6C**). Specifically, miR-24, miR-9, miR-125b, miR-21, miR-221, miR-17, miR-1, and miR-30 emerged as regulators in *mid-stage* hESC-RPE-sEVs, whereas miR-30 and miR-141 are present as regulators in *late-stage* hESC-RPE-sEV (**Figure 6D**). KEGG molecular interaction maps based on miRNA target genes retrieved from the IPA analysis output show the implication of mid-stage hESC-RPE-sEV-associated miRNAs in PEDF signaling, specifically highlighting their involvement in the regulation of apoptosis, neuroprotection, cell cycle, and differentiation (**Figure 6D**).

**Figure 6.**
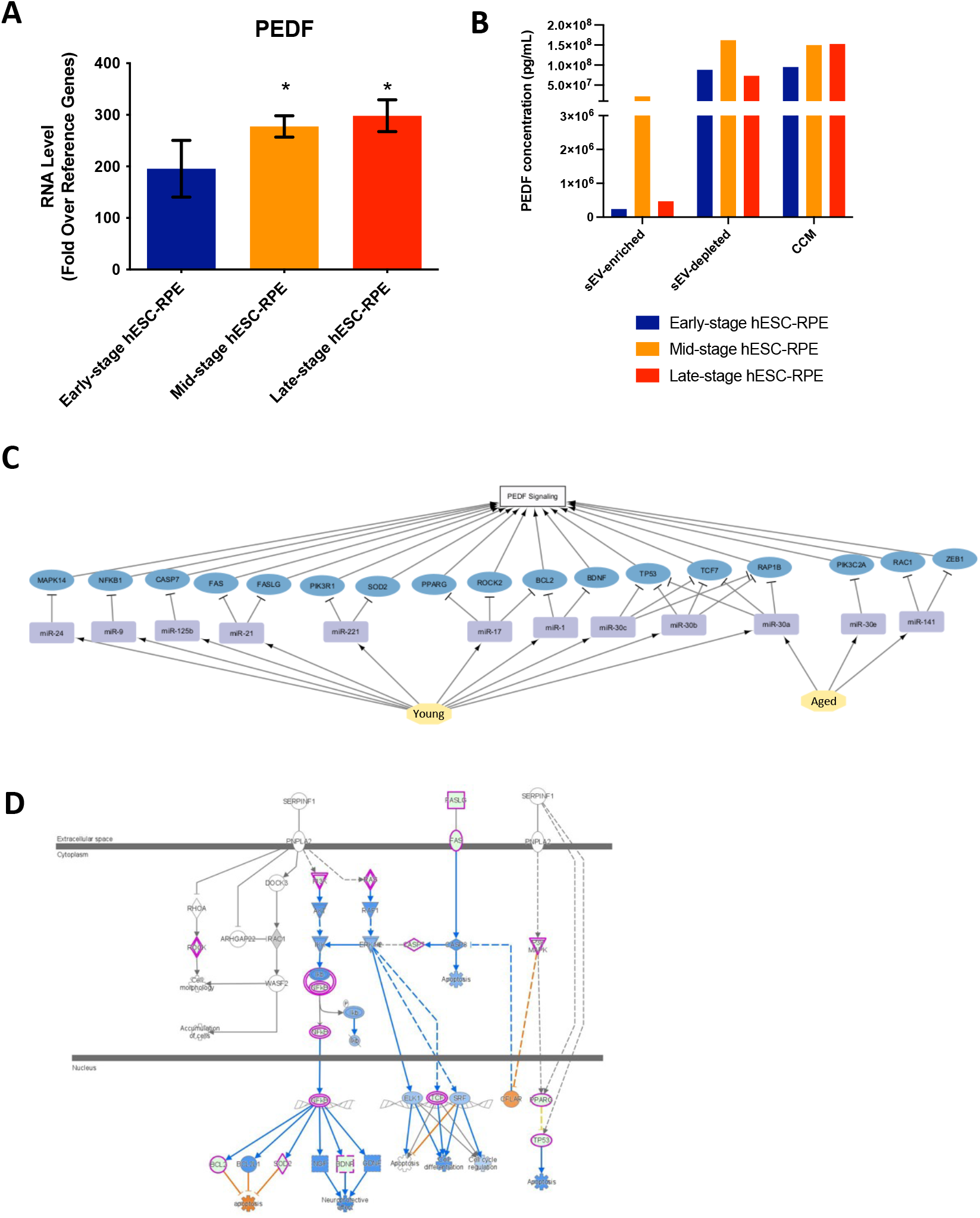
**A.** The gene expression of pigment epithelium-derived factor (PEDF) in hESC-RPE cells gradually increased during maturation in hESC-RPE-sEVs across the *early-stage* (11-22 days in culture)*, mid-stage* (28-39 days in culture), and *late-stage* (59-70 days in culture) (*N*=4). **B.** PEDF secretory function of the hESC-RPE cells was evaluated by ELISA on the conditioned cell culture media (CCM) and in two fractions of the hESC-RPE secretome: sEV-fraction and sEV-depleted (CCM minus sEV). Results indicated a progressive increase in PEDF concentration in CCM during the maturation of hESC-RPE. The sEV-depleted fraction maintained consistent PEDF levels across all groups while the sEV-enriched fraction exhibited the highest PEDF concentration in the *Mid-stage* hESC-RPE group. **C.** Networks exploring age-dependent miRNA-gene targets-disrupted PEDF-related pathway interactions using Cytoscape. **D.** KEGG Molecular Interaction Map showing PEDF signaling pathways associated with miRNA in *mid-stage* hESC-RPE-sEV. * *p* < 0.05

## Discussion

Several previous studies have reported variability in cellular miRNA expression during the differentiation of RPE from stem cells, as well as the involvement of different target miRNAs from the RPE in mediating retinal diseases (Wooff et al., 2020; Ahn et al., 2021; Morris et al., 2020; Guo et al., 2023). **19, 34-37** However, to the best of our knowledge, our study is the first to conduct a comprehensive analysis of the miRNA profile contained in EVs secreted from hESC-RPE during their maturation. As stem cell-derived RPE cell therapy advances towards clinical application, research has revealed the in vitro cellular maturation of stem cell-derived RPE, which can extend up to 8 weeks in culture. Interestingly, their therapeutic efficacy in vivo appears to be better in cells cultured for 4 weeks (Schwartz et al., 2015; Mehat et al., 2018; Mandai et al., 2017; Kashani et al., 2018; Kashani et al., 2021; Al-Ani et al.,2020; Davis et al., 2017). **X** However, there is a lack of studies focusing on determining the optimal phenotype of EV-secreting RPE and understanding the biological cargo of EVs to develop RPE-EV-based therapeutics.

Our study results reveal an interesting finding that the number of recovered sEV from *mid-stage* hESC-RPE, while maintaining similar cell culture numbers and the same volume input of CCM, was significantly higher compared to early-stage and late-stage hESC-RPE. This observation suggests a potentially biologically active status of cells in terms of sEV secretion during *mid-stage* maturation (28-39 days in culture), which aligns with peak therapeutic efficacy in hESC-RPE cell transplantation therapy, as opposed to earlier or later stages (Davis et al., 2017). X Notably, our previous research demonstrated higher EV particle numbers in vitreous samples from proliferative vitreoretinopathy (PVR) patients compared to non-PVR patients, supporting numbers of EV secretion is regulated either in physiologic or pathologic conditions (Nair et al.,2022). **40** Despite differences in particle numbers, other characterizations of EVs, including size, morphology, and comprehensive EV profiles of the three groups of hESC-RPE-sEV, were comparable. Regarding miRNA profiling, the heat map and pie chart illustrate the distinctly different types of miRNAs and levels of expression of secreted miRNAs via sEV from hESC-RPE at different maturation statuses. The overall miRNA expression pool consisting of 94 miRNAs (*Ct* values ≤ 30) contained in hESC-RPE-sEV displayed varying counts of miRNA in sEV secreted from *early-, mid-, and late-stage* hESC-RPE cells, totaling 27, 85, and 34 miRNAs, respectively. This indicate that *mid-stage* hESC-RPE-sEV contain a greater variety of miRNAs compared to hESC-RPE of other stages, as well as a higher number of secreted sEV. It is noteworthy that no miRNA was exclusively expressed in *early-* or *late-stage* hESC-RPE-sEV, suggesting that the miRNA profiling reflects the two extreme ends of the spectrum during the maturation of the parental cells. Nevertheless, it can not be determined from the current dataset whether the increased secretion of miRNA is directly correlated with the increased numbers of secreted sEV. Furthermore, it is unknown whether all sEV in a population contain the same amount or same kind of microRNAs or if some vesicles are selectively loaded with specific microRNAs, given the heterogeneity of cell-secreted sEV. Despite these limitations stemming from current technological constraints, our study results provide valuable information on the increased numbers of secreted sEVs and greater variety of miRNAs contained in *mid-stage* hESC-RPE-sEVs. This result is validated by rigorous characterization of the RPE cell phenotypes and secreted sEV using our established workflow (Leung et al, 2024). After recovering sEV from CCM, we performed comprehensive orthogonal biophysical and biomolecular characterizations of hESC-RPE-sEVs to accurately define their profile. Additionally, we assessed the purity of the recovered sEV, including evaluations of total protein, particle-to-protein ratio, and total RNA concentration of the hESC-RPE-sEV. Furthermore, the bead-based multiplex flow cytometry assay (MACSPlex) analysis in hESC-RPE was distinctly different from sEV secreted by HEK 293, serving as a nonocular cell control. This suggests that our comprehensive approach to characterize sEV was sensitive enough to distinguish sEV from different cell sources.

In the pathway analysis to determine the molecular pathways involved in these differentially expressed miRNAs at each stage of hESC-RPE-sEV, it is not surprising that we found more pathways associated with miRNAs in sEV secreted from *mid-stage* hESC-RPE. Out of the total 511 different pathways identified from the total pool of 94 miRNAs, our focused analysis on pathways related to “EVs” or “retina” revealed several significant pathways. Specifically, EV trafficking, tissue and cellular polarity, growth factor regulation, inflammation, and oxidative stress & senescence emerged as major pathways associated with EVs and retina. miRNA contained in sEV from *early-stage* hESC-RPE were predominantly associated with EV trafficking and RPE structures such as clathrin endocytosis and tissue polarity and epithelial junction. Meanwhile, miRNA contained in sEV from *mid-stage* and *late-stage* hESC-RPE were more broadly involved in these five major pathways. Despite ongoing human clinical trials using hESC-RPE cells with similar cell phenotypes (cultured for around 4 weeks) based on the protective therapeutic efficacy observed in pre-clinical studies, the current study cannot verify whether these pathways are protective or detrimental to retinal health (e.g., in inflammation or oxidative stress) (Ke et al., 2021; Bian et al., 2020; Mathew et al., 2019). Further studies, including investigations into the therapeutic efficacy and potential toxicity of RPE-sEV treatment in retina disease models, as well as efforts to determine key protective microRNAs, are warranted for future research endeavors. Based on our study results, it would be a logical choice to prioritize testing mid-stage hESC-RPE-sEV initially.

In addition, our study results highlight that the miRNA contents in sEV from hESC-RPE differs substantially from the cellular miRNA content observed in previous studies supporting that cellular miRNA and EV mediated miRNA are differently regulated (Yuan et al, 2015). While the precise functional role of miRNA contained in hES-RPE-sEV requires further exploration in specific signaling contexts, our pathway analyses support at least three potential roles for miRNA in hESC-RPE-sEV. Firstly, they may be involved in EV machinery playing as EV resident miRNAs, such as EV trafficking, wherein three main mechanisms for EV-RNA uptake: cell surface membrane fusion, cell contact-dependent mechanisms, and endocytic pathways, such as clathrin-mediated, caveolin-dependent, receptor-mediated endocytosis, micropinocytosis, and phagocytosis. Secondly, these miRNAs may contribute to RPE cell health, impacting tissue and cellular polarity. This suggests that secreted miRNA indeed may mediate the microenvironment and/or health of RPE, possibly through autocrine and/or paracrine effects. Thirdly, miRNAs might play a role in multiple pathways associated with essential RPE functions, such as growth factor regulation, inflammation, and oxidative stress. Consequently, miRNAs in hESC-RPE-sEV could participate in both EV machinery-related processes and functional roles of the RPE.

With the excitement surrounding the abundance of miRNA associated with essential RPE functions contained in RPE-sEV from our study results, the key question regarding the therapeutic relevance of EV miRNA remains their capacity to initiate phenotypic effects in recipient cells through cytoplasmic delivery. Joshi et al. and Bonsergent et al. reported that approximately 10% to 30% of internalized EVs released cargo to the cytoplasm, as indicated by their reporter system (Joshi et al., 2020; Bonsergent et al., 2021). Jong et al. observed that the functional transfer of RNA varied based on the combination of donor and receptor cells (de Jong, 2020). Additionally, it was established that EV-delivered miRNAs could participate in mRNA suppression (Abels et al., 2019). Therefore, there is great therapeutic potential for miRNA contained in EV. Along the way, we identified additional challenges due to the limited tools available to define and predict the multidimensional synergistic benefits from EV therapy when multiple molecules simultaneously affect the microenvironment. We observed that pigment epithelium-derived factor (PEDF), a major trophic factor of RPE, was expressed in a substantial portion of the soluble factor compartment and a smaller pool of sEV. Interestingly, miRNAs, including miR-24, miR-9, miR-125b, miR-21, miR-221, miR-17, miR-1, miR-30, and miR-141, were found to be involved in PEDF signaling pathways. We believe that these results are from a controlled cargo sorting mechanism rather than coincidence, supporting the possibility that multimolecular (protein and miRNA)-based sEV therapeutics may offer a synergistic benefit to the biological system. Next-generation integrative pathway enrichment analysis of multivariate omics data is necessary for this gap.

This study has limitations. The assessment was conducted under uniform cell culture conditions. However, it is still possible that in vivo RPE-secreted sEV have different miRNA profiling reflecting the microenvironment. While defining the optimal in vitro cell culture conditions to produce optimal miRNA-containing sEV in order to develop RPE-sEV therapeutics is a necessary step, it would be additionally helpful to investigate sEV-contained miRNA profiling from RPE exposed to different cell conditions. Furthermore, our study results are specific to hESC-derived RPE. Another prevalent source, iPSC-derived RPE, will require a separate investigation.

In summary, the microRNAs found in hESC-RPE-sEVs reveal their potential diverse roles, from aiding in vesicle movement to supporting cell health and participating in essential RPE cellular functions. The differences between vesicle miRNA content from our study and cellular miRNA from the previous study highlight the unique way these vesicles sort their cargo. While our study provides important insights into identifying the optimal phenotype of the EV secreting cell, more research is needed to fully understand how these microRNAs work and their potential effects. Exploring these complexities could lead to innovative treatments for various diseases, making vesicle-delivered microRNAs a promising area for future therapies for retinal diseases.

## Material and Methods

### hESC-RPE Cell Differentiation, and Culture

Human embryonic stem cells (WA09, WiCell Research Institute) were spontaneously differentiated into RPE cells as previously described (Croze et al., 2014). Briefly, colonies of hESCs were manually passaged, seeded onto tissue culture plates coated with Matrigel hESC-Qualified Matrix (Corning #354277), and received biweekly medium exchanges of serum free XVIVO-10 medium (Lonza #(BE) BP04-743Q) for approximately three months. Pigmented regions were then manually isolated, expanded for two passages, and cryopreserved at 2-5 days post-seed as a cellular suspension using CryoStor10 cryopreservation medium (BioLife Solutions #210102). Thawed cells were expanded using 10uM Y-27632 (Tocris #1254) until the eighth passage as previously described (Leach et al., 2016). Cultures were tested every other month using MycoAlert Mycoplasma Detection Kit (Lonza LT07-318) and determined to be negative for mycoplasma throughout the study.

### Pigmentation Assessment

Upon reaching confluence, for each age group, representative fields of view were acquired by bright field and phase contrast microscopy using an Olympus CK40 inverted microscope and Infinity Capture software. Upon reaching confluence, pigmentation in hESC-RPE cells was categorized as follows: grade 0 – pigmentation in 0-10% of the total area, grade 1 – pigmentation in 11-25% of the total area, grade 2 – pigmentation in 26-50% of the total area, grade 3 – pigmentation in 50-80% of the total area, and grade 4 – pigmentation in >80% of the total area (Bennis et al., 2007; Ahluwalia et al., 2023).

### HEK293T Cell Culture

As for HEK293T cell culture, 2 x 10^6^ cells were cultured in 10 mL high glucose Dulbecco’s Modified Eagle’s Medium (Thermo Fisher Scientific, Waltham, MA) containing sodium pyruvate (110 mg/L), L-glutamine (200 mm), 10% fetal bovine serum and 1% penicillin (10,000 U/mL) /streptomycin (10,000 µg/mL) in a 10 cm culture dish at 37°C in a 5% CO_2_. HEK293T cells at their tenth passage were used in this study.

### Immunocytochemistry and Confocal Fluorescence Microscopy

hESC-RPE were cultured on sterile, matrigel-coated, #1.5 coverslips (GG121.5PRE, Neuvitro) before fixation with 4% methanol-free formaldehyde in PBS for 20 min. Cells were subsequently permeabilized with 0.1% Triton X-100 for 10 min, blocked with 5% goat serum (Jackson ImmunoResearch) and 1% BSA (Thermo Fisher Scientific) in PBS for 30 min, and incubated in the following primary antibodies overnight at 4°C: rabbit-anti-ZO1 (40-2200, Thermo Fisher Scientific, 1:200) and mouse-anti-RPE65 (MAB5428, Millipore-Sigma, 1:200). Coverslips were washed three times with PBS followed by secondary antibody incubation in blocking buffer for 1 hr at room temperature. Secondary antibodies were: AlexaFluor 594 AffiniPure goat anti-rabbit IgG (111585144, Jackson ImmunoResearch, 1:200) and AlexaFluor 488 AffiniPure goat anti-mouse IgG (115545062, Jackson ImmunoResearch, 1:200). Nuclei were stained for 10 min with Hoechst 33342 in PBS (2 μg/mL), washed three times with PBS, mounted in ProLong Gold antifade, and imaged on an Olympus FV1000 Spectral Confocal with a PLAPON-SC 60X oil objective (NA: 1.40), and excitation laser lines at 405, 488, 559, and 635 nm. ImageJ-FIJI (NIH, USA) was used to generate single confocal planes extracted from z-stacks.

### RT-qPCR of hESC-RPE Cells

Lysates for RNA purification were prepared by aspirating growth medium and triturating hESC-RPE cultures in RLT Buffer (Qiagen) followed by storage at -80°C. RNA was purified using RNeasy columns (Qiagen) and included an on-column genomic DNA digestion step using RNase-free DNase (Qiagen). Reverse transcription and PCR were performed on 35ng of total RNA per well using AgPath-ID One-Step RT-PCR Reagents (4387424, Thermo Fisher Scientific) with the following TaqMan primer-probe sets (all Thermo Fisher Scientific): EIF2B2 (Hs00204540_m1), SERF2 (Hs00428481_m1), UBE2R2 (Hs00215107_m1), BEST1 (Hs00188249_m1), RPE65 (Hs01071462_m1), and PEDF (Hs01106934_m1). All Ct data were first linearized (2^-Ct^), and values for RPE65, BEST1, and PEDF were normalized to the geometric mean of the three reference genes (EIF2B2, SERF2, and UBE2R2).

### Preparation of Conditioned Cell Culture Medium

To generate conditioned cell culture medium (CCM) from hESC-RPE evaluation, RPE cells were enzymatically passaged using TrypLE Select (Gibco #12563011) per manufacturer’s instructions and seeded onto tissue culture plates coated with Matrigel hESC-Qualified Matrix (Corning #354277) at a density of 70,000 cells/cm^2^. The viability of the cells at the time of plating was determined by trypan blue exclusion, and cultures exceeding 95% viability were used for the study. At one day post-seed, cells were rinsed with DPBS (Gibco #14040141) and fed with 4mL XVIVO-10 medium (Lonza #(BE)BP04-743Q) per well within a 6-well tissue culture plate. The medium was exchanged twice per week and was supplemented with 10mM Y-27632 (Tocris #1254) for 10-15 days until the RPE cells attained cuboidal morphology. Starting at 11 until 70 days post-seed, CCM was collected from four wells of a 6-well plate using a serological pipet, combined, and immediately frozen in a 50mL Falcon centrifuge tube at -80°C until future use. Four replicate plates were used at each collection time point. The incubation period in which cells conditioned the medium ranged from 3-4 days at 37°C, 5% CO_2_. The three different passage time ranges were selected based on RPE pigmentation and cell proliferation speed while maintaining other RPE cell characteristics including RPE65, BEST1, and PEDF expression, with days of gap between the groups to avoid any potential molecular overlapping (Dunn et al., 1996; German et al., 2008; Bennis et al., 2007; Al-Ani et al.,2020; Davis et al., 2017).

To generate conditioned CCM from HEK293T cells, once the cell confluency reached 90% the culture medium was changed to 10 mL XVIVO-10 medium (Lonza #(BE)BP04-743Q) containing 1% penicillin (10,000 U/mL) /streptomycin (10,000 µg/mL) and maintained for another 4 days at 37°C in a 5% CO_2_. Collected conditioned CCM was immediately frozen in a 50mL Falcon centrifuge tube at -80°C until future use.

### Small Extracellular Vesicle Recovery

For sEV recovery, ExoDisc® (LabSpinner, Ulsan, South Korea), a microfluidic tangential flow filtration method-based equipment was used following the manufacturer’s instructions (Woo et al., 2017; Dong et al., 2020; Leung et al., 2024). In brief, an input volume of 3 mL of each test article (early-stage hESC-RPE, mid-stage hESC-RPE, late-stage hESC-RPE, and HEK293T) was processed on an ExoDisc® using the bench-top operating machine (OPR-1000, LabSpinner™, South Korea). Purified sEV were retrieved from the collection chamber using 100 μL of PBS and immediately stored at −80 °C until further use within 2-4 weeks.

### Biophysical Particle Analysis

The concentration and size distribution of recovered sEV were measured by ZetaView (Particle Metrix, Germany) following the manufacturer’s protocol. In brief, ZetaView (NV) analysis, samples were diluted to achieve an ideal particle per frame value of 150–200 particles/frame. For each measurement, three cycles were performed by scanning 11 cell positions each and capturing 60 frames per position under the following settings: autofocus, camera sensitivity for all samples: 85, shutter: 70, and cell temperature: 25°C. Following the data capture, the videos were subjected to analysis using the built-in ZetaView Software version 8.02.31 (Leung et al., 2024).

### Transmission Electron Microscopy

sEV were visualized by negative-stained transmission electron microscopy (TEM) using a JEOL JEM-2100 microscope mounted on a Gatan OneView IS camera. Thin formvar/carbon-coated EM grids (Ted Pella, Inc.) were loaded with 6 µL diluted sEV solution, incubated for 4 minutes, excess solution wicked, and stained with 10 µL 4% uranyl acetate for 3 minutes. After staining, the excess solution was removed, and the grid was allowed to dry for 10 minutes before storage for future TEM observation at 80 kV.

### Single-Particle Interferometric Reflectance Imaging Sensing: ExoView Analysis

Single Particle Interferometric Reflectance Imaging Sensing (SP-IRIS) with the ExoView R100 system and the ExoView Human Tetraspanin Kit (NanoView Biosciences, USA) was used (Leung et al., 2024). Each testing article was incubated on an ExoView Tetraspanin Chip for 16 hours at room temperature, followed by three washes in solution A (ExoView Human Tetraspanin Kit, NanoView Biosciences, USA). Pre-diluted immunocapture antibodies (anti-CD9 CF488, anti-CD81 CF555, and anti-CD63 CF647) were used at a 1:500 dilution in solution A. To achieve the correct antibody concentration, 250 microliters of the antibody solution were mixed with the remaining 250 μL of solution A after chip washing, resulting in a final antibody dilution of 1:1000 for incubation. After a 1-hour incubation at room temperature, the chips were washed, dried, and imaged using the ExoView R100 reader and ExoView Scanner 3.0 acquisition software, and the data were subsequently analyzed using the ExoView Analyzer 3.0.

### Bead-Based Multiplex Flow Cytometry Assay (MACSPlex) Analysis

A MACSPlex human Exosome kit (Miltenyi Biotec, Bergisch-Gladbach, Germany) was used, following the manufacturer’s protocol, to assess the expanded surface protein markers of sEV (Leung et al., 2024). The sEV were captured using 37 distinct surface marker antibodies (CD1c, CD2, CD3, CD4, CD8, CD9, CD11c, CD14, CD19, CD20, CD24, CD25, CD29, CD31, CD40, CD41b, CD42a, CD44, CD45, CD49e, CD56, CD62p, CD63, CD69, CD81, CD86, CD105, CD133.1, CD142, CD146, CD209, CD326, HLA-ABC, HLA-DR DP DQ, MCSP, ROR1 and SSEA-4) simultaneously and include the two isotype controls (mIgG1 and REA control) corresponding to the antibodies conjugated with fluorescent beads and then analyzed via flow cytometry. The processed samples were run on a Cytek Aurora Flow Cytometer (Cytek Biosciences, USA) and analyzed with SpectroFlo software (Cytek Biosciences, USA).

### Protein Concentration Analysis

The recovered sEV were lysed by mixing 5 μL of each testing article with 5 μL of 2x RIPA buffer. Protein concentrations of the lysed sEV preparations were quantified using the Pierce™ BCA Protein Assay Kit (Thermo Fisher Scientific, USA), following the manufacturer’s protocols (Leung et al., 2024).

### PEDF ELISA

The concentration of PEDF concentration was assessed using a human PEDF ELISA kit (Abcam #ab246535). Various fractions of *Juvenile*, *Young*, and *Aged* hESC-RPE-derived CCM, including the sEV-enriched fraction, sEV-depleted fraction, and total unprocessed CCM were subjected to analysis. ELISA procedures were conducted following the manufacturer’s protocol. The absorbances were measured immediately at 450 nm in a microplate reader (SpectraMax iD5, Molecular Devices, CA, U.S.A.). For standardization, the PEDF concentration was normalized by the protein concentration of the samples.

### Total RNA Extraction and miRNA Quantification

The total RNA extraction from sEV was performed using an exoRNeasy midi kit (QIAGEN, Hilden, Germany) according to the manufacturer’s protocol (Leung et al., 2024). In brief, pre-isolated sEV were mixed with an equal volume of a binding buffer (XBP) and added to the membrane affinity column. After discarding the flow-through using a centrifuge, washing buffer (XWP) was added to the column; and the sEV were mixed with a lysis reagent (QIAzol®) passing through the membrane and extracted to the bottom of the tube. Then, chloroform was added to remove the phenol component; and the clear layer containing the isolated RNA was separated through centrifugation. After several washing steps, the total RNA was finally obtained. The purity and quantity of the isolated total RNA were measured using a Nanodrop™ 2000 spectrophotometer (Thermo Fisher Scientific, Wilmington, DE).

### microRNA Profiling

TaqMan® MicroRNA Reverse Transcription Kit and the Megaplex™ RT Primers were used to synthesize single-stranded cDNA from total RNA recovered from the testing sEV samples according to the manufacturer’s instructions. Briefly, 4.5 μL of the RT Reaction Mix was added to designated wells in a reaction plate or tubes, along with 2 μL of RNA or water (no template control), ensuring a total RNA content of 2 ng. The samples underwent a 5-minute incubation on ice, followed by thermal cycling with standard ramp speed and a reaction volume of 7.5 μL. Following reverse transcription, the obtained cDNA underwent preamplification using Megaplex™ PreAmp Primers. Specifically, 2.5 μL of the cDNA was added to wells of a reaction plate along with 22.5 μL of TaqMan® PreAmp Master Mix and incubated using a standard ramp speed following the manufacturer’s settings. Post-amplification, the reaction plate was briefly centrifuged, and 75 μL of 0.1X TE pH 8.0 was added to each well. The diluted pre-amplified products were then utilized directly for real-time PCR reactions according to the manufacturer’s protocol (Applied Biosystems™ TaqMan™ Array Human MicroRNA A+B Cards Set v3.0, ThermoFisher Scientific). For the data analysis, the Ct value of an endogenous control gene (U6) was subtracted from the corresponding Ct value for the target gene resulting in the ΔCt value which was used for relative quantification of miRNA expression. As there is an inverse correlation between ΔCt and miRNA expression level, lower ΔCt values were associated with increased miRNA expression. Since a Ct value of 30 represents single molecule template detection, Ct values > 30 were considered to be below the detection level of the assay. Therefore, only the miRNAs with a Ct ≤ 30 were included in the analyses. The DDCt method was applied to calculate the relative expression levels (fold change) of the target miRNAs. Fold-change values of the selected miRNAs were transformed to log2 values. Differential miRNA expressions were determined by performing a two-sample t-test for each probe.

### Bioinformatics and Pathway Analysis

Differentially expressed miRNAs were analyzed, and their experimentally confirmed gene targets from the miRBase database, supported by literature references, were retrieved using The Ingenuity Knowledge Base as a reference set. Additional information on significant miRNAs and their target genes was obtained using IREDESCENT, a proprietary software scanning MEDLINE. Target gene sets for each age group’s significant miRNAs were combined and studied for overrepresentation in documented functional gene sets. Ingenuity Pathway Analysis (IPA) identified impacted pathways, regulatory interactions, and networks, with KEGG molecular interaction maps retrieved for significant pathways. Cytoscape was used to construct and visualize age-dependent miRNA-gene target-disrupted pathway interactions, while R-studio functions were employed to generate heatmaps displaying miRNA overlapping pathways for each group of statistically significant pathways.

### Statistical Analysis

Unless specified otherwise, data are presented as mean ± standard deviation (SD). Statistical analysis and graph plotting were performed using GraphPad Prism. Student’s t-test was utilized to compare the two groups, and significance was defined as *p-value* < 0.05.

## Acknowledgments

The authors thank NRI-MCDB Microscopy Facility Spectral Laser Scanning Confocal (award S10OD010610). The authors thank Extracellular Vesicle Core at Children’s Hospital Los Angeles and Paolo Neviani, PhD for their technical assistance with ExoView analysis.

---

**Supplementary Figure 1.**
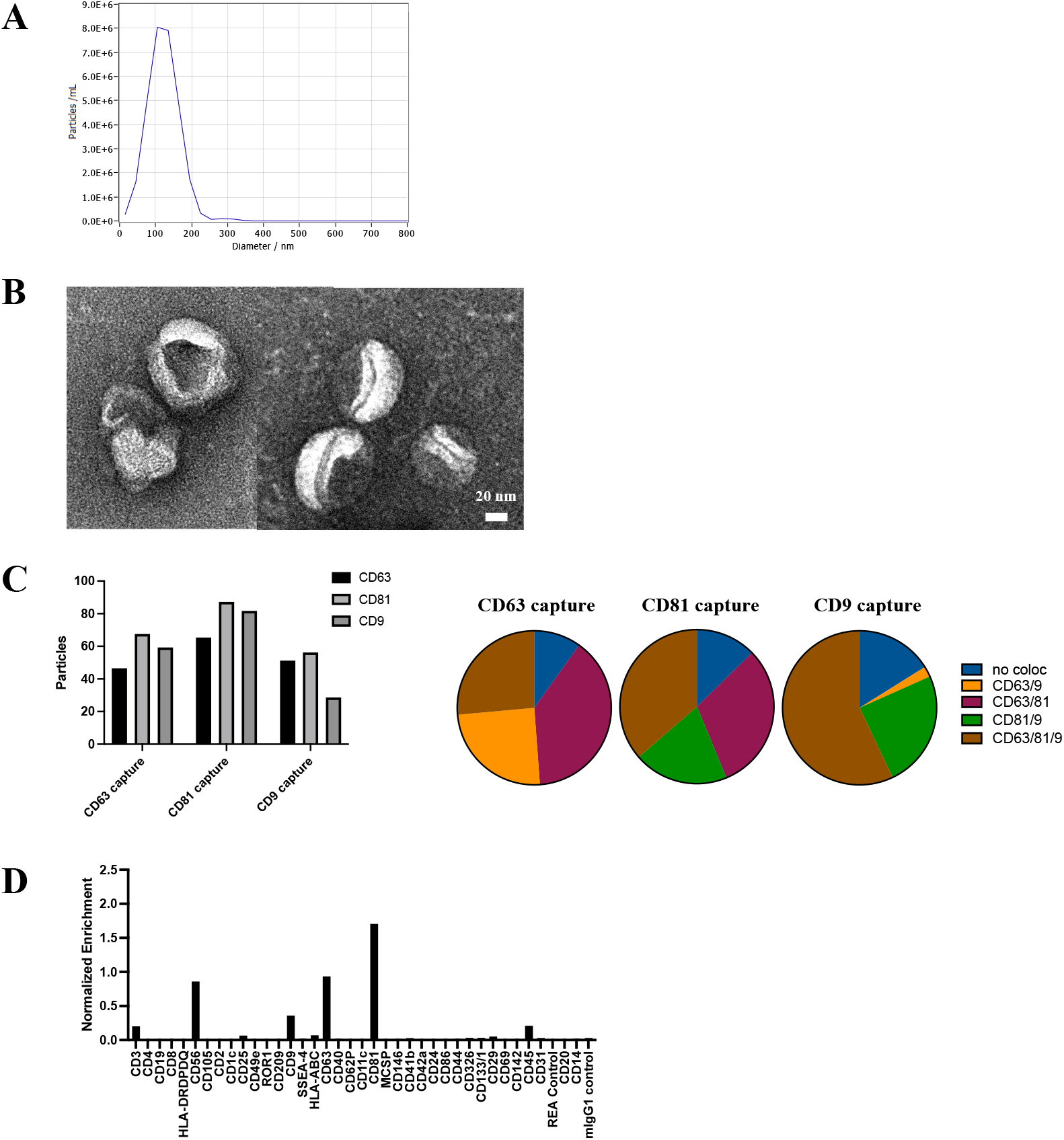
HEK293T cell-derived sEV characterization based on: **A.** NTA (ZetaView), **B.** TEM, **C**. ExoView analysis, and **D.** MACSPlex analysis.

**Supplementary Figure 2.**
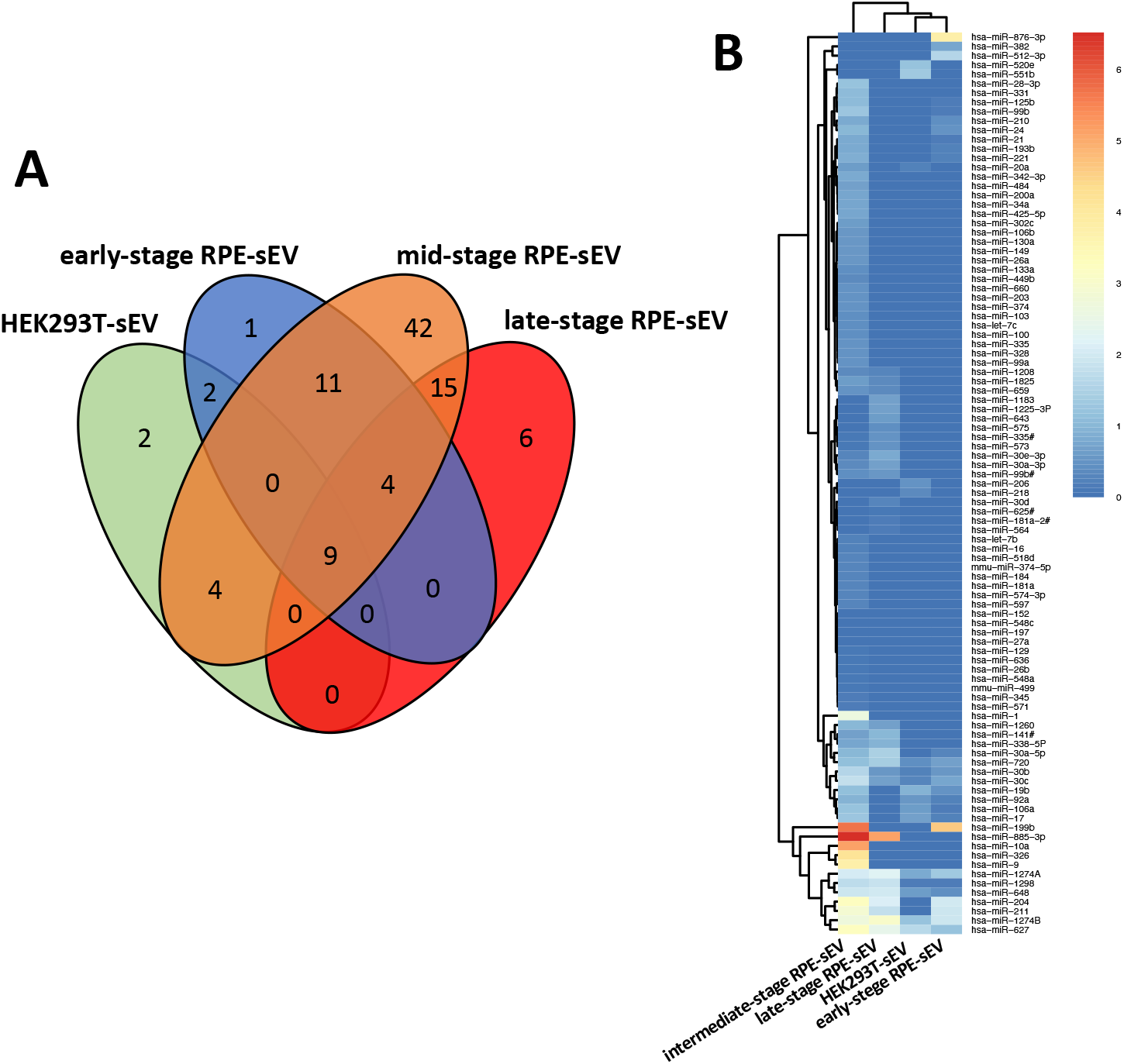
**A.** Venn diagram of the total number of miRNAs identified in the sEV derived from the three hESC-RPE and the HEK293T cell groups. **B.** Heatmap of all the expressed miRNAs in hESC-RPE- and HEK293T-sEV.

**Supplementary Figure 3.**
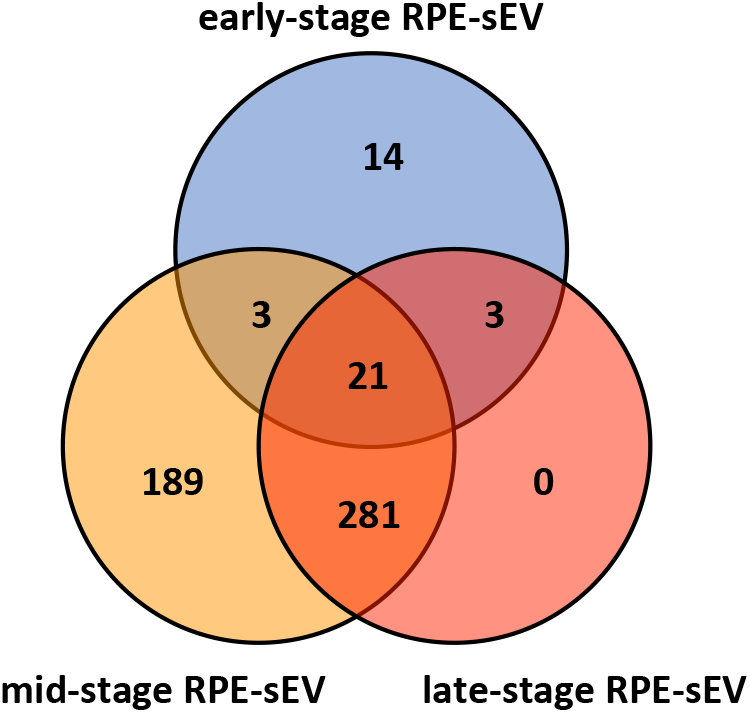
Venn diagram of the total 511 statistically significant pathways associated with the miRNAs expressed in hESC-RPE-sEV identified by Ingenuity Pathway Analysis (IPA).

**Supplementary Table 1.** miRNA expression in *early-stage* hESC-RPE-sEV.

| Early-stage hESC-RPE-sEV |  |
| --- | --- |
| Target Name | fold-change |
| hsa-miR-199b | 39031.54 |
| hsa-miR-876-3p | 5776.59 |
| hsa-miR-204 | 97.42 |
| hsa-miR-1274B | 84.31 |
| hsa-miR-211 | 80.61 |
| hsa-miR-512-3p | 37.41 |
| hsa-miR-1274A | 20.17 |
| hsa-miR-627 | 14.13 |
| hsa-miR-30c | 4.88 |
| hsa-miR-382 | 4.50 |
| hsa-miR-720 | 3.93 |
| hsa-miR-30b | 2.56 |
| hsa-miR-19b | 2.19 |
| hsa-miR-24 | 2.02 |
| hsa-miR-210 | 1.77 |
| hsa-miR-648 | 1.67 |
| hsa-miR-30a-5p | 0.83 |
| hsa-miR-221 | 0.68 |
| hsa-miR-17 | 0.62 |
| hsa-miR-193b | 0.60 |
| hsa-miR-92a | 0.58 |
| hsa-miR-1298 | 0.52 |
| hsa-miR-106a | 0.51 |
| hsa-miR-125b | 0.45 |
| hsa-miR-21 | 0.40 |
| hsa-miR-99b | 0.39 |
| hsa-miR-130a | 0.02 |

**Supplementary Table 2.** miRNA expression in *mid-stage* hESC-RPE-sEV.

| Mid-stage hESC-RPE-sEV |  |
| --- | --- |
| Target Name | fold-change |
| hsa-miR-885-3p | 3275297.93 |
| hsa-miR-199b | 499886.49 |
| hsa-miR-10a | 132479.71 |
| hsa-miR-326 | 13278.00 |
| hsa-miR-9 | 5516.34 |
| hsa-miR-204 | 1795.30 |
| hsa-miR-627 | 1651.96 |
| hsa-miR-211 | 744.05 |
| hsa-miR-1274B | 494.63 |
| hsa-miR-1 | 408.35 |
| hsa-miR-1274A | 120.12 |
| hsa-miR-648 | 79.38 |
| hsa-miR-30c | 56.47 |
| hsa-miR-1298 | 49.67 |
| hsa-miR-30b | 38.10 |
| hsa-miR-17 | 16.51 |
| hsa-miR-99b | 16.36 |
| hsa-miR-106a | 16.19 |
| hsa-miR-28-3p | 14.15 |
| hsa-miR-19b | 13.77 |
| hsa-miR-720 | 12.07 |
| hsa-miR-331 | 11.54 |
| hsa-miR-125b | 11.30 |
| hsa-miR-24 | 9.94 |
| hsa-miR-1260 | 9.79 |
| hsa-miR-30a-5p | 9.53 |
| hsa-miR-92a | 8.18 |
| hsa-miR-221 | 6.68 |
| hsa-miR-193b | 5.68 |
| hsa-miR-210 | 5.60 |
| hsa-miR-342-3p | 5.50 |
| hsa-miR-21 | 5.21 |
| hsa-miR-425-5p | 4.75 |
| hsa-miR-34a | 4.74 |
| hsa-miR-200a | 4.50 |
| hsa-miR-338-5P | 4.38 |
| hsa-miR-484 | 4.17 |
| hsa-miR-141 | 3.86 |
| hsa-miR-20a | 3.46 |
| hsa-miR-1825 | 3.28 |
| hsa-miR-302c | 3.06 |
| hsa-miR-106b | 2.79 |
| hsa-miR-26a | 2.68 |
| hsa-miR-149 | 2.64 |
| hsa-miR-130a | 2.59 |
| hsa-miR-660 | 2.31 |
| hsa-miR-374 | 2.24 |
| hsa-miR-203 | 2.21 |
| hsa-miR-659 | 2.20 |
| hsa-miR-335 | 2.07 |
| hsa-miR-328 | 1.93 |
| hsa-miR-99a | 1.90 |
| hsa-miR-100 | 1.82 |
| hsa-let-7c | 1.77 |
| hsa-miR-103 | 1.71 |
| hsa-miR-133a | 1.58 |
| hsa-miR-99b# | 1.46 |
| hsa-miR-449b | 1.37 |
| hsa-miR-1208 | 1.20 |
| hsa-miR-30a-3p | 1.12 |
| hsa-miR-518d | 1.09 |
| mmu-miR-374-5p | 1.08 |
| hsa-miR-30e-3p | 0.99 |
| hsa-miR-181a | 0.97 |
| hsa-miR-597 | 0.94 |
| hsa-miR-574-3p | 0.94 |
| hsa-miR-184 | 0.88 |
| hsa-let-7b | 0.77 |
| hsa-miR-16 | 0.62 |
| hsa-miR-345 | 0.40 |
| hsa-miR-571 | 0.39 |
| mmu-miR-499 | 0.33 |
| hsa-miR-573 | 0.32 |
| hsa-miR-636 | 0.27 |
| hsa-miR-26b | 0.25 |
| hsa-miR-548a | 0.24 |
| hsa-miR-335 | 0.22 |
| hsa-miR-129 | 0.19 |
| hsa-miR-1225-3P | 0.15 |
| hsa-miR-218 | 0.13 |
| hsa-miR-152 | 0.06 |
| hsa-miR-548c | 0.05 |
| hsa-miR-27a | 0.03 |
| hsa-miR-197 | 0.02 |
| hsa-miR-643 | 0.01 |

**Supplementary Table 3.**
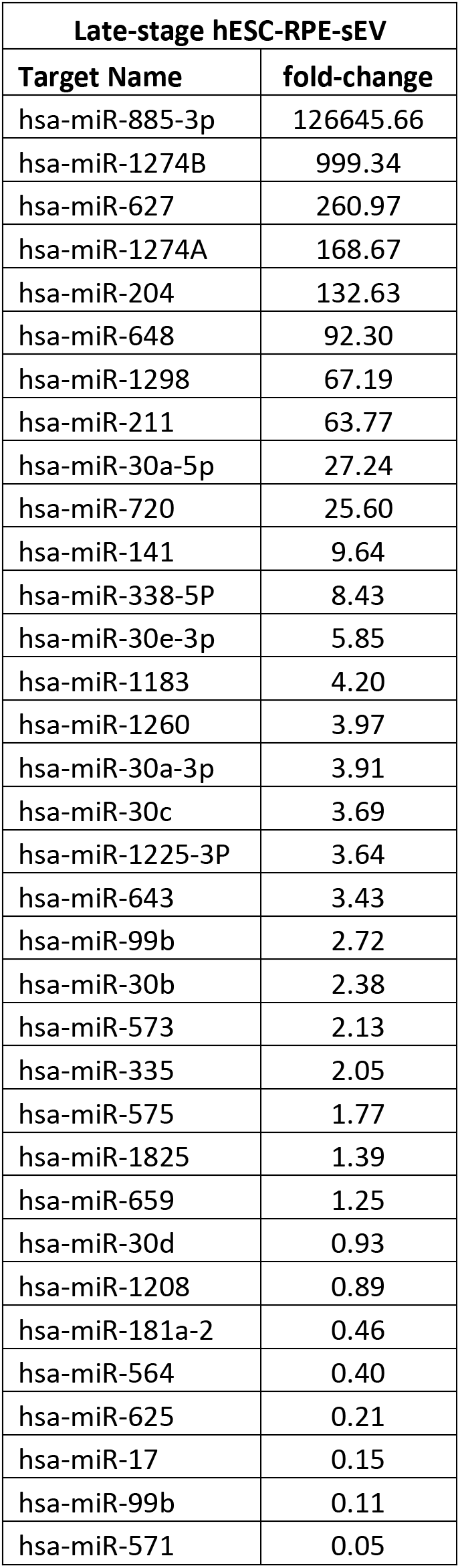
miRNA expression in *late-stage* hESC-RPE-sEV.

| Late-stage hESC-RPE-sEV |  |
| --- | --- |
| Target Name | fold-change |
| hsa-miR-885-3p | 126645.66 |
| hsa-miR-1274B | 999.34 |
| hsa-miR-627 | 260.97 |
| hsa-miR-1274A | 168.67 |
| hsa-miR-204 | 132.63 |
| hsa-miR-648 | 92.30 |
| hsa-miR-1298 | 67.19 |
| hsa-miR-211 | 63.77 |
| hsa-miR-30a-5p | 27.24 |
| hsa-miR-720 | 25.60 |
| hsa-miR-141 | 9.64 |
| hsa-miR-338-5P | 8.43 |
| hsa-miR-30e-3p | 5.85 |
| hsa-miR-1183 | 4.20 |
| hsa-miR-1260 | 3.97 |
| hsa-miR-30a-3p | 3.91 |
| hsa-miR-30c | 3.69 |
| hsa-miR-1225-3P | 3.64 |
| hsa-miR-643 | 3.43 |
| hsa-miR-99b | 2.72 |
| hsa-miR-30b | 2.38 |
| hsa-miR-573 | 2.13 |
| hsa-miR-335 | 2.05 |
| hsa-miR-575 | 1.77 |
| hsa-miR-1825 | 1.39 |
| hsa-miR-659 | 1.25 |
| hsa-miR-30d | 0.93 |
| hsa-miR-1208 | 0.89 |
| hsa-miR-181a-2 | 0.46 |
| hsa-miR-564 | 0.40 |
| hsa-miR-625 | 0.21 |
| hsa-miR-17 | 0.15 |
| hsa-miR-99b | 0.11 |
| hsa-miR-571 | 0.05 |

**Supplementary Table 4.** miRNA expression in HEK293T-sEV.

| HEK293T-sEV |  |
| --- | --- |
| Target Name | fold-change |
| hsa-miR-106a | 3.28 |
| hsa-miR-1274A | 5.58 |
| hsa-miR-1274B | 14.47 |
| hsa-miR-1298 | 0.53 |
| hsa-miR-17 | 3.52 |
| hsa-miR-19b | 8.00 |
| hsa-miR-206 | 2.13 |
| hsa-miR-20a | 0.66 |
| hsa-miR-218 | 1.34 |
| hsa-miR-30b | 0.77 |
| hsa-miR-30c | 0.86 |
| hsa-miR-520e | 16.20 |
| hsa-miR-551b | 20.16 |
| hsa-miR-627 | 46.86 |
| hsa-miR-648 | 3.33 |
| hsa-miR-720 | 1.51 |
| hsa-miR-92a | 2.46 |

